# Integrating XMALab and DeepLabCut for high-throughput XROMM

**DOI:** 10.1101/2020.04.10.035949

**Authors:** JD Laurence-Chasen, AR Manafzadeh, NG Hatsopoulos, CF Ross, FI Arce-McShane

## Abstract

Marker tracking is a major bottleneck in studies involving X-ray Reconstruction of Moving Morphology (XROMM). Here, we tested whether DeepLabCut, a new deep learning package built for markerless tracking, could be applied to videoradiographic data to improve data processing throughput. Our novel workflow integrates XMALab, the existing XROMM marker tracking software, and DeepLabCut while retaining each program’s utility. XMALab is used for generating training datasets, error correction, and 3D reconstruction, whereas the majority of marker tracking is transferred to DeepLabCut for automatic batch processing. In the two case studies that involved an *in vivo* behavior, our workflow achieved a 6 to 13-fold increase in data throughput. In the third case study, which involved an acyclic, *post mortem* manipulation, DeepLabCut struggled to generalize to the range of novel poses and did not surpass the throughput of XMALab alone. Deployed in the proper context, this new workflow facilitates large scale XROMM studies that were previously precluded by software constraints.

## INTRODUCTION

Data processing in kinematics workflows can be a time-consuming and laborious task, especially when three-dimensional (3D) reconstruction requires the integration of data from multiple cameras. In marker-based XROMM (X-ray Reconstruction of Moving Morphology; Brainerd et al., 2010), every radiopaque marker in every frame of two X-ray videos must be accurately tracked. This step has been streamlined by the open-source program XMALab (Knörlein et al., 2016), which offers a suite of features for marker detection, visualization, and tracking. Marker tracking remains a major bottleneck in the XROMM workflow, however, limiting the feasibility of studies that require large sample sizes across multiple individuals or species (cf. Gintof et al., 2010; Granatosky et al., 2019; Iriarte-Diaz et al., 2017; Martinez et al., 2018).

In the past several years, deep learning, a type of machine learning, has emerged as a powerful tool for automating pose estimation in kinematics workflows (Graving et al., 2019; Insafutdinov et al., 2016; Mathis and Mathis, 2019, for a recent review; Pereira et al., 2019). In particular, the open-source deep learning toolbox DeepLabCut (Mathis et al., 2018; Nath et al., 2018) has been rapidly and widely adopted in the scientific community. DeepLabCut was designed for markerless, automatic tracking of body parts in light camera videos and has been used in a disparate range of study systems with impressive performance and robustness (Labuguen et al., 2019; Owen et al., 2019; Parmiani et al., 2019; Stringer et al., 2019, and many others).

The degree to which DeepLabCut’s utility in digitizing light videos can be transferred to the biplanar videoradiographic data at the core of XROMM is not known. The properties of videoradiographic data render marker tracking a challenging process to automate. Whereas in light video different body parts are immediately distinguishable by their shape and appearance alone, in X-ray videos all markers can be identical in appearance (small black spheres), and thus only identifiable in their broader spatiotemporal context. Moreover, as many XROMM studies aim to quantify subtle motions, the desired error threshold in marker tracking is extremely small (i.e., ≤1 pixel; Brainerd et al., 2010). The graphical user interface (GUI) and reconstruction features of XMALab are specifically designed for the accurate identification and tracking of markers under these challenging conditions.

The purpose of this paper is to present a workflow that integrates DeepLabCut into the existing XROMM data processing pipeline—retaining the XMALab labeling GUI and reconstruction tools while offloading initial batch tracking to DeepLabCut. We compare the performance of our pipeline to the standard XMALab workflow on three different datasets, each with different behaviors and marker sets. Strengths and weakness of the two different pipelines are discussed, and instructions and recommendations for the full implementation of our pipeline are provided.

## MATERIALS AND METHODS Software Availability

Open-source Python code under the name XROMM_DLCTools and Jupyter Notebooks for the full implementation of our integrated workflow are available at https://github.com/jdlaurence/xromm_dlctools. We used DeepLabCut version 2.1 for all analysis and testing (github.com/AlexEMG/DeepLabCut).

### Data flow

The flow of data through our pipeline is cyclic (Fig. 1). A training dataset—a relatively small subset of paired camera-1 and camera-2 frames—is tracked as accurately as possible in XMALab. Then those tracked data, in the form of 2D points and their corresponding images, are migrated to DeepLabCut, where they are used to train an artificial neural network. Different videos can then be directly input to DeepLabCut for automated tracking. After the network predicts the marker locations in the new videos, the predicted 2D points are brought back into XMALab for error correction, performance evaluation, and 3D reconstruction. If the network’s performance is sub-optimal then areas of high error/poor performance can be manually corrected in XMALab, those corrected frames added to the training dataset, and the process repeated until the desired performance is achieved.

**Figure 1.**
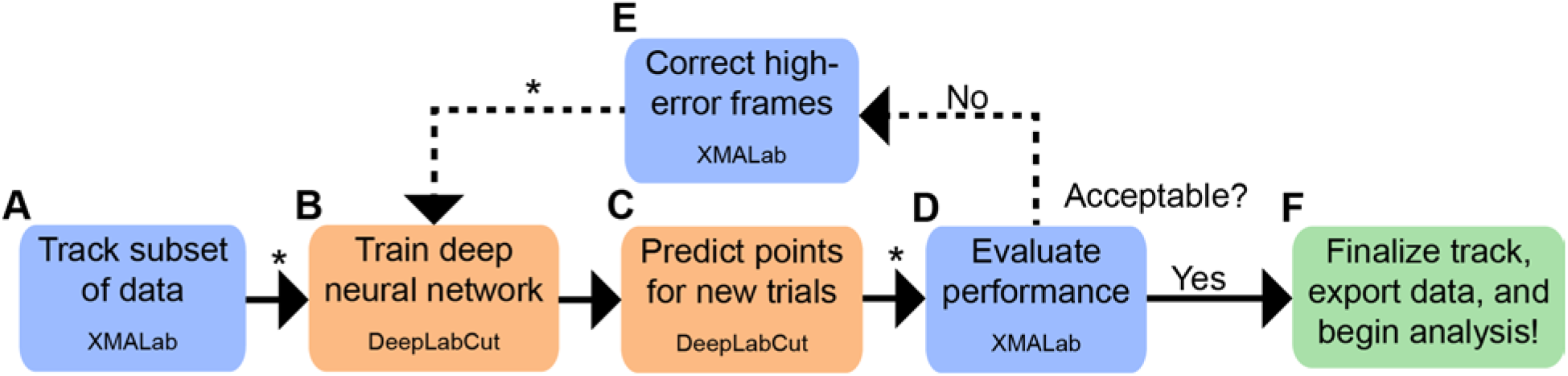
Integrated XMALab – DeepLabCut workflow for marker tracking. (A) The user begins by tracking 200-500 frames from the dataset in XMALab. (B) Those frames serve as the training dataset for a deep neural network trained with DeepLabCut. (C) That network can then predict 2D points for new trials. (D) The predicted points are imported back into XMALab and measures of tracking quality (e.g., reprojection and rigid body error) are used to determine whether the project-specific performance criteria are met. (E) If errors are too high, select frames can be corrected, added to the training dataset, and steps B-D repeated. (F) Once performance is acceptable, the user corrects any remaining errors in XMALab, and can export the data (3D points and rigid body transformations) for analysis. (*) indicates the step is performed by an XROMM_DLCTools function.

### Training dataset generation

The composition of the training dataset is perhaps the single most important factor in DeepLabCut’s performance. The network will generalize, i.e., perform well on new trials, when frames in the training dataset completely capture the variation in the full dataset. In other words, a pattern-recognizing neural network is liable to fail when faced with totally novel patterns, in this case marker postures. Thus, for each test case, we tracked 250-500 consecutive frames from 3-6 trials (approx. 800-2000 frames total), carefully selecting those regions that contained the most postures and/or stages of the behavior of interest. Once all frames were tracked and 2D points exported from XMALab, we created a DeepLabCut project using its built-in methods. The project configuration file was edited to match the specifics of the dataset (i.e., marker names, file location paths, etc.).

After the project was successfully created, we used the DLCTools Python function *xma_to_dlc* to create a DeepLabCut-ready training dataset. The function reads XMALab 2D points files and extracts data from frames with tracked data. It also extracts the frame of video itself, either from an avi file, or jpg stack, and converts it to a png image. The function loops through the provided trials, picking one frame at a time until the total number of picked frames equals the ‘nframes’ argument, or until all tracked frames have been picked. The output of the function is identical to the output of the native DeepLabCut labeling GUI; thus after this step, the user proceeds to the established DeepLabCut workflow, starting with the function *create_training_dataset*.

Given the redundancy of postures inherent in consecutive frames of high-speed video, as well as the added computational cost of a larger training dataset, we uniformly subsampled the tracked frames by setting the ‘nframes’ argument to either 500 or 750. This meant the training dataset was composed of every other initially tracked frame. The choice to create an initial training dataset with more frames than the recommended number for a DeepLabCut study (~200) was made based on the inherent visual complexity and challenge of identifying multiple, visually homogeneous markers in X-ray videos. The impact of smaller and larger initial training datasets on performance is discussed in the results section.

### Network training and analysis

Once the training dataset is generated, the standard DeepLabCut workflow is followed. The functions *create_training_dataset* and *train_network* were used to train a single neural network whose weights were optimized for both camera-1 and camera-2 videos. In all cases, we used ResNet-101. We allowed training to run until DeepLabCut’s native cross-entropy loss function plateaued, typically between 200,000 and 500,000 iterations. DLCTools supports the use of separate neural networks for each camera plane, if the user chooses. This would double the amount of training but may improve performance. The DLCTools function *analyze_xromm_videos* calls the native *analyze_videos* function to predict points for new trials. It automatically detects the camera-1 and camera-2 videos and combines DeepLabCut’s predicted points output into a single 2D file in XMALab format. The predicted 2D points files can then be imported into a XMALab file with the corresponding calibration.

### Performance evaluation

We evaluated the trained neural network’s tracking performance in two ways. The first method, which we do not recommend to be used exclusively, consists of DeepLabCut’s evaluation function, which measures the mean 2D distance between the predicted points and the user-provided (via XMALab) ‘ground-truth’ points for the test fraction of the training dataset. In our experience, this native *evaluate_network* function is not a sufficient measure of tracking quality for XROMM data for several reasons: 1) Marker-specific error values are not provided, 2) camera calibration information is not used to measure 3D error (see Discussion), 3) the function can be affected by over-fitting of the network to the training dataset, and 4) the function cannot, by definition, assess the performance of a network’s tracking of a novel trial. In other words, the error values provided by *evaluate_network* may not indicate that the network is ready to generalize and perform adequately on novel trials.

The second means of evaluating the network’s tracking performance, which we recommend, is the suite of error measurement tools in XMALab. This uses individual marker reprojection error (see Knörlein et al., 2016) as an overall heuristic for tracking performance. As the goal of marker tracking with DeepLabCut is to accelerate the process while maintaining accuracy, we determined the reprojection error value at which the measured variables did not significantly differ by tracking mode. These variables were joint coordinate system (JCS; Grood and Suntay, 1983) data and, in the case of the tongue data set, 3D marker positions.

For each network iteration (see following section), we tested a novel trial that had also been tracked in XMALab alone. Thus, there were two sets of data for that trial: data tracked with DeepLabCut, and data tracked with XMALab. We took the tracked data through the XROMM pipeline and then calculated the mean difference between corresponding variables across all test frames. We deemed the tracking acceptable if this value is smaller than the precision threshold for that variable (*sensu* Menegaz et al., 2015). When this threshold was reached, the reprojection error values for all points were recorded, as was the time spent for post-DeepLabCut corrections in XMALab.

This approach necessitates rigorous tracking in XMALab for the training dataset and for the basis of comparison (i.e., precision study). When evaluating DeepLabCut output, single frames or regions of frames that exhibit critically poor tracking may not be captured by the mean reprojection or rigid body error. Thus, it is *essential* that the predicted points data be migrated into XMALab and the reprojection error, 2D position, and rigid body error plots be visually inspected for large outliers.

### Training dataset augmentation

When DeepLabCut’s tracking quality is not satisfactory, areas of poor performance can be manually corrected and added to the training dataset for the network to ‘learn’. The user identifies high-error frames by visual inspection of the reprojection error and rigid body error traces, as well as the gestalt-appearance of the tracking in the main window; i.e., are the crosshairs on the markers? The exact numbers depend on the specific study, but typically involve reprojection errors over 2 pixels, and rigid body errors over 0.05 cm. Once the user corrects all markers in that frame, they add the frame number to a frame index spreadsheet that contains the trial name and frames corrected from that trial. The DLCTools function *add_frames* reads this file, extracts the corrected frame images and their new 2D point data, and appends them to the training dataset. The user can then repeat network training and re-analyze the same videos with improved marker prediction. Exact file format and folder structure for the use of this function are detailed in the online package instructions.

### Test cases

In order to assess the accuracy and limitations of our new workflow, we tested it on three previously collected datasets: pig feeding, monkey feeding, and bird leg range of motion (ROM). Importantly, the datasets share few similarities; they were collected on three different biplanar radiography systems and differ in species, number of markers, marker size, and marker locations. Example images from each dataset are provided (Supp. Fig. 1). While these case studies are certainly not exhaustive in terms of taxa or behaviors, their differences provide a basis for evaluating the degree to which this workflow can be generalized to future XROMM studies.

For each case study, we report the training parameters, reprojection error values, and time spent digitizing to achieve reconstructions that are statistically indistinguishable from those made from data tracked in XMALab alone (following the methods described above). As correction in XMALab is the final step in both workflows, and subject to user bias, we did not perform additional inferential statistics across conditions (e.g., mean reprojection error). The variables used to make this comparison are detailed in the following sections. For all cases, ResNet-101 was used and training was stopped when the cross-entropy loss plateaued or fell below 0.005, typically at 200,000-500,000 iterations.

### Study 1: Minipig Feeding

These publicly available minipig (*Sus scrofa*) feeding data were collected with C-arm videofluoroscopes, and have been used in XROMM tutorials and software testing for the last decade (Brainerd et al., 2010; Knörlein et al., 2016). Ten 1 mm tantalum markers—five in the cranium and five in the mandible—exhibit typical difficult-to-track characteristics; as the pig feeds unconstrained, the markers occasionally cross and occlude one another, or enter areas of low contrast. The first iteration of the training dataset comprised a total of 500 frames from three trials of SusD feeding (dataset 2006-12-29). The network was then tested on a novel trial—specifically, the 435-frame trial from the same date that has been used for previous teaching and validation studies. We used the six degrees of freedom from the temporomandibular JCS as the output variables for performance comparisons.

### Study 2: Macaque Feeding

In this study, performed at The University of Chicago XROMM Facility, a rhesus macaque (*Macaca mulatta*) fed on grapes and gummy bears while head-fixed. The data were collected at 200 Hz with an X-ray technique of 100-105 kilovolt peak (kVp) and 10-12.5 milliamperes (mA). A total of 24 tantalum markers, all 1 mm in diameter, were located as follows: 4 in the cranium, 4 in the mandible, 1 in the hyoid, and 15 in the tongue. The tongue markers, being in a soft body, moved in complex ways, frequently crossing and occluding one another. The combination of numerous bone and soft tissue markers makes these data extremely difficult to track; an expert XMALab user took approximately 8 hours to track a 10 second, 2000 frame trial. The first iteration of the training dataset comprised 750 frames sampled from six trials. Following sub-optimal performance on the test trial, the training dataset was augmented twice, such that the final training dataset comprised 1250 frames. Final manual corrections of output involved setting a reprojection error threshold at 2-2.5 pixels and correcting all frames that exceeded that threshold. We used the temporomandibular joint rotation values and tongue point 3D positions as the output variables for performance comparisons.

### Study 3: Guineafowl Range of Motion

In this study, performed at the W.M. Keck Foundation XROMM Facility at Brown University, the hind limb of a helmeted guineafowl (*Numida meleagris*) was physically manipulated, *post mortem*, to assess the ROM of the bird’s hip, knee, and ankle joints. The data were collected at 50 Hz with an X-ray technique of 70-85 kVp and 200 mA. A total of 12 0.8 mm zirconium oxide markers were placed in the pelvis, femur, tibiotarsus, and tarsometatarsus–3 in each element. Unlike the previous two studies, these data do not involve a cyclic behavior; in fact, the aim of the study was to explore each joint’s full ROM through intentionally non-cyclic, non-repeated movements (Kambic et al., 2017; Manafzadeh and Padian, 2018). An expert XMALab user took approximately 10 hours to track an 1800 frame trial. The first iteration of the training dataset comprised 750 frames from four trials. The training dataset was augmented twice, and the final training dataset comprised 1500 frames. We used the rotations at the three joints as the output variables for performance comparisons.

## RESULTS

### Study 1: Minipig Feeding

When applied to the pig feeding dataset, DeepLabCut rapidly reached XMALab-level performance. Cross-entropy error during network training plateaued at approximately 250,000 iterations. DeepLabCut’s raw (i.e. pre-XMALab correction) marker predictions for a novel trial exhibited rigid body errors and JCS rotation values that fell within the precision threshold of the study (Fig. 2A,C). Mean reprojection error of individual points, however, was higher in the trial tracked with DeepLabCut (0.51 ± 0.25 s.d. pixels) as compared with XMALab (0.16 ± 0.02 pixels). This difference in mean reprojection error persisted after manual correction of select, high error frames in XMALab. Nevertheless, the difference in measured JCS variables never fell outside of the error threshold. DeepLabCut was immediately robust to the cyclic crossing of select markers that consistently required manual intervention when tracking in XMALab. After training the neural network on the initial training dataset, time to fully track 1000 frames decreased from approximately 30 minutes with XMALab alone to 5 minutes with the integrated workflow, constituting a six-fold increase in throughput when tracking cranial and mandibular markers.

**Figure 2.**
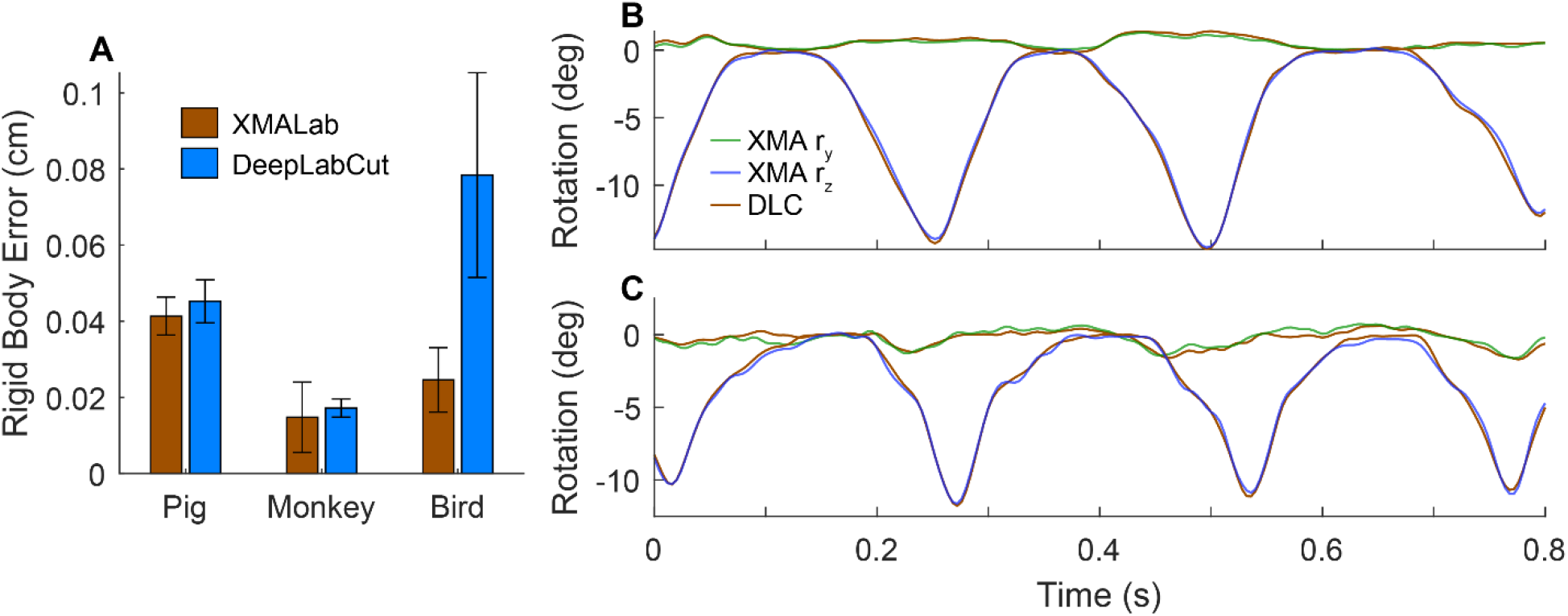
Comparison of XMALab and DeepLabCut rigid body tracking performance. (A) Mean rigid body error (filtered at 30 Hz) from XMALab for the test trial of each case study where markers were tracked either with XMALab (brown) or DeepLabCut (blue). Here, the DeepLabCut marker predictions were not corrected in XMALab. Pig and monkey errors correspond to the rigid body transformations (cranium and mandible) used to animate and extract temporomandibular joint rotation data from a subset of the monkey (B) and pig (C) test trials. The bird errors are the mean of all leg bones for the test trial. Note that despite differing mean reprojection errors (see main text), mean rigid body error and resultant rotation values for the pig and monkey were comparable to that of the same trial tracked in XMALab. Joint coordinate systems were oriented following Menegaz et al. (2015) and Orsbon et al. (2018). The rx (x-axis rotation) trace was omitted because it failed to exceed the established noise threshold in both tracking methods.

### Study 2: Macaque Feeding

The visually challenging nature of these data was the original motivation for the present study. Complex tongue deformations during chewing result in a wide range of marker crossings and occlusions. Moreover, maintaining consistent marker identity with a large amount of intermarker motion is difficult. These challenges were reflected in DeepLabCut’s tracking; DeepLabCut rapidly achieved XMALab-level performance when tracking the markers in the two rigid bodies—the cranium and mandible. Before any manual correction, mean rigid body error was comparable to that of the trial tracked using XMALab only. Likewise, the temporomandibular JCS Y- and Z-axis rotations fell within the respective variable’s precision thresholds (Fig. 2A,B). As in the pig dataset, DeepLabCut-predicted marker locations exhibited higher mean reprojection errors, both before and after manual correction, than the XMALab trial.

For every iteration of the network, the uncorrected XYZ positions of the tongue markers did not meet the threshold for successful performance (Fig. 3A-D). The first iteration of the network produced predictions that required approximately two hours of manual correction per 2000 frame trial to reach the error threshold. Satisfactory performance was achieved through the correction of all frames in which a marker’s reprojection error exceeded 2-2.5 pixels (Fig. 3E-F). In order to reduce the amount of manual correction needed, new frames were tracked and added to the training dataset two separate times. In each case, the cross-entropy loss plateaued at approximately 600,000 iterations of network training. The output of the second iteration of the network required one hour of manual correction, and the third iteration of the network required twenty to thirty minutes, an approximately 13-fold increase in throughput, including training dataset generation time. We stopped network re-training after the second augmentation of the training dataset, although it is possible that the amount of per-trial manual correction could be further reduced with additional augmentations.

**Figure 3.**
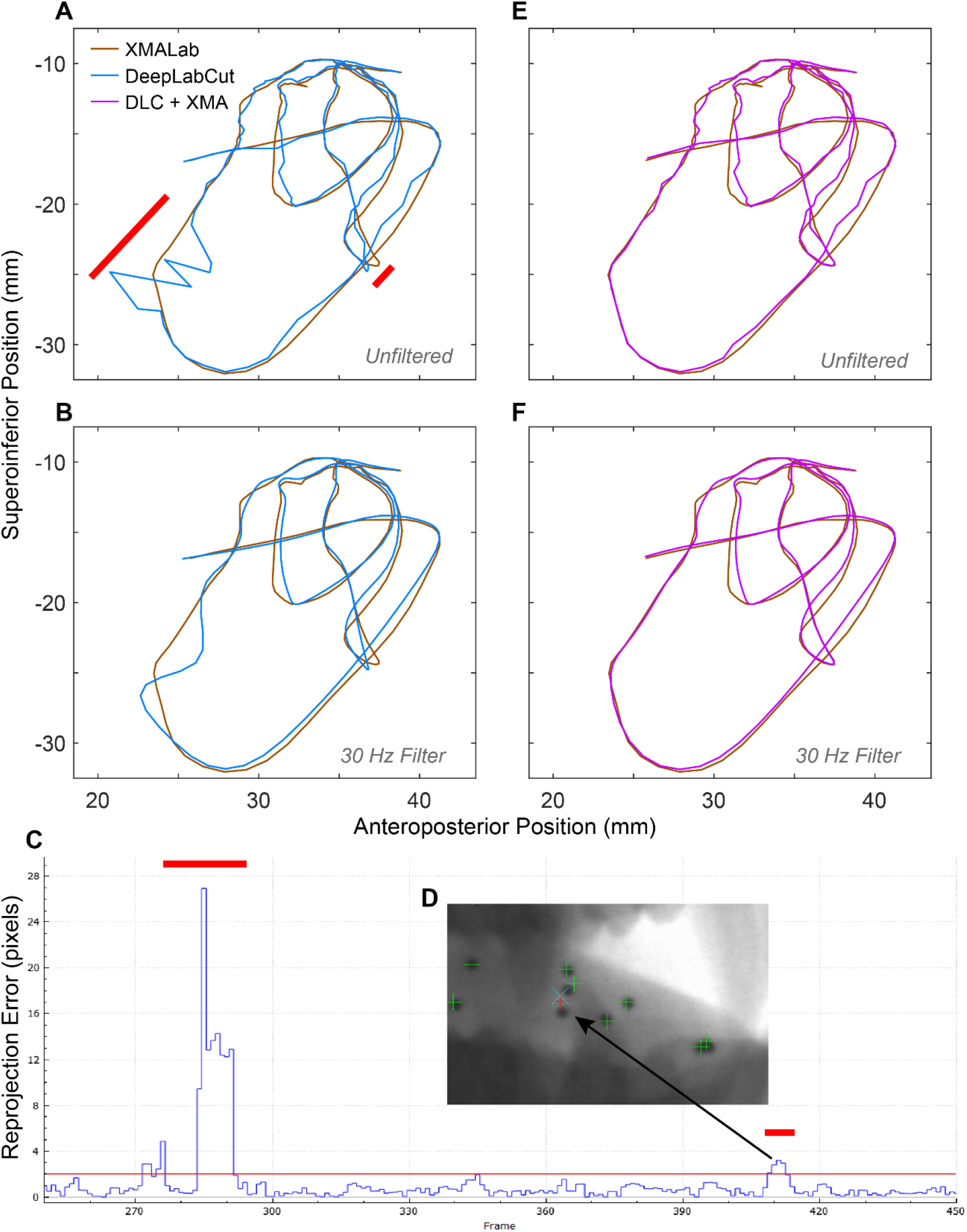
Comparison of tracking methods for an example tongue marker trajectory. (A-D) DeepLabCut’s predicted positions (blue line) for the anterior right tongue marker were at times erroenous (red bars). After importing the predicted 2D points into XMALab, those regions of poor tracking were easily identified with the reprojection error trace (C). All frames with reprojection errors higher than the established threshold (2 pixels for this study) were corrected in both cameras (D). (E-F) The resulting DeepLabCut + XMALab marker trajectory (magenta line) is accurate, and 8-13x faster to generate. The trajectories are the X and Y values taken from the marker’s XYZ coordinates that have been exported from XMALab, and transformed into an anatomical coordinate system with its origin at the posterior nasal spine (Orsbon et al. (2018). Unfiltered (A & E) and filtered with a 30 Hz low-pass butterworth filter (B & F) trajectories are shown.

### Study 3: Guineafowl Range of Motion

This case study was unique in that the “behavior” being studied—*post mortem* specimen manipulation—was acyclic and designed to document the range of possible poses. In short, we were unable to achieve successful marker tracking results with DeepLabCut. After three iterative augmentations of the training dataset, reprojection and rigid body errors were still so high that it took longer to correct the output of DeepLabCut than to track the test trial from scratch in XMALab (Fig. 2A). As each trial contained novel postures, we found it was virtually impossible to generate a training dataset that sufficiently captured the variation in the data without tracking a majority of every trial in XMALab—defeating the purpose of the new workflow.

## DISCUSSION

### Comparison with XMALab

In two of the three case studies, our integrated workflow dramatically outperformed XMALab alone, in terms of overall processing time. After network optimization, per-trial marker tracking time was reduced 6-fold in the pig dataset and 13-fold in the monkey dataset. This high-throughput performance was robust to marker placement and number; the monkey dataset involved >20 markers in both rigid bodies (cranium and mandible) and soft tissue (tongue) structures. At the individual point level, DeepLabCut converged on, but never surpassed, XMALab quality. We found that mean reprojection errors of individual points were lowest when tracked in XMALab alone, but, crucially, this difference was not reflected in measured output variables. After correction in XMALab, both JCS data and XYZ marker positions did not differ significantly between the two tracking modes. We interpret the elevated reprojection errors in DeepLabCut to be the result of greater noise, much of which can be easily filtered. In general, we found that what was difficult in XMALab was also more difficult for DeepLabCut; whereas DeepLabCut excelled at tracking markers in rigid bodies that followed cyclic trajectories, it had more difficulty (i.e., required more training frames) for dense and overlapping markers in soft tissue.

### Establishing an error tolerance

Here, we set the error tolerance for marker tracking in DeepLabCut as the reprojection error and rigid body error values that corresponded to the point at which the measured variable (JCS data, or tongue marker positions) did not differ significantly from the same variable when tracked in XMALab alone. Depending on the nature of the study at hand, different performance criteria may be desired. For example, if a study is constrained to a small number of trials, noise inherent to DeepLabCut’s predictions can have a magnified impact and thus more stringent error tolerances are appropriate. Likewise, in a study that seeks to quantify subtle motions (e.g., hemimandible wiggle, Bhullar et al., 2019), extra care must be taken when establishing the error tolerance.

### Training dataset

One of the most important factors that influence the performance of a deep neural network is the degree to which the training dataset captures the range of variation in the broader dataset to be analyzed (Tan et al., 2018). Thus, algorithmic selection of training data based on visual dissimilarity can greatly improve performance for a given training dataset size (Brust et al., 2019). DeepLabCut offers a *k*-means method for extracting frames from videos that show maximum visual differences. In theory, this approach could be used on XROMM data, however, in practice it is generally not feasible as it can be virtually impossible to accurately identify markers in single frames of XROMM data out of their temporal context. For this reason, the workflow involves tracking sub-sequences of trials and it is up to the user to identify the regions that contain different postures. In the future, an algorithmic approach to identify ideal training frames could reduce time spent augmenting the training dataset.

### Other factors influencing throughput

We found that markers in rigid bodies were much more consistently tracked by DeepLabCut. This likely due to the fact that the relative positions of the markers stay consistent and the bones themselves are constrained by their articulations. In other words, the postural space of markers in bones is much more constrained than a set of markers in soft tissues like the tongue, and thus easier to sample for a training dataset.

Image resolution has a dramatic impact on DeepLabCut processing time (Mathis et al., 2018). As such, best practice is to down-sample large images before processing. We chose to omit any image down-sampling due to the small size of XROMM markers (<10 pixels diameter) and our desire to maximize performance. For studies where markers are larger or processing time is of greater concern, down-sampling the raw X-ray data may yield better results. The hardware on which a user runs DeepLabCut should also be considered. Without a dedicated GPU, processing full-sized XROMM images becomes practically infeasible.

The extent to which a user asks the network to generalize is also a major factor influencing throughput. In the traditional DeepLabCut workflow, networks are meant to be generalized across individuals and days of data collection. Here, we performed data analysis on a single individual from each case study as different individuals had a different marker locations and marker numbers. Whether or not a single network can be generalized across multiple days of data collection probably depends on the variation in the day-to-day setup, as well as the marker locations.

### Areas for improvement

XMALab and DeepLabCut utilize fundamentally different mechanisms for marker tracking. XMALab uses a point’s velocity to make a prediction about where it will be in the following frame, then searches for the point using a template. Additionally, it uses camera calibration information like reprojection error and rigid body error for user visualization as well as to establish thresholds at which to stop tracking. In contrast, DeepLabCut uses neither camera calibration information nor velocity when tracking. DeepLabCut evaluates each frame of video in isolation, essentially pattern-matching the appearance of the frame at-hand to frames from the training dataset. The lack of communication between the two camera views means that DeepLabCut might make a highly-erroneous prediction when, to a user looking at both cameras views simultaneously, it is obvious that marker correspondence is incorrect. If DeepLabCut employed a filter based on reprojection error (see DLTdv8; Hedrick, 2008) and used a marker’s velocity, tracking performance might improve dramatically.

### Concluding Remarks

We showed that a marker tracking workflow that integrates deep learning can dramatically outperform the existing XROMM workflow in terms of throughput. The increased speed did not impact the accuracy of measured kinematics variables, though reprojection errors in XMALab were higher. Importantly, the throughput increase occurred when the behavior at hand was cyclic and when ROM was constrained experimentally. For this reason, we believe it is best to think about the present workflow as one that enables large scale studies, the likes of which were previously impossible when such experimental design criteria are met. This workflow is not, however, a cure-all for digitizing XROMM data. In cases where the sample size is small, or the behavior is acyclic, the established XMALab-only marker tracking workflow is still more efficient. As deep learning algorithms improve, however, and when DeepLabCut incorporates camera calibration into its marker prediction, this balance will likely shift.

## ACKNOWLEDGEMENTS

We thank Greg Shakhnarovich and Steven Basart for helpful discussions on using deep neural networks for XROMM data processing, Ben Knorlein for helpful discussions on integrating DeepLabCut with XMALab, Alexander Mathis and Mackenzie Mathis for DeepLabCut, and David Baier for XROMM_MayaTools. Rebecca Junod provided superb assistance with monkey data collection. We also thank Victoria Hosack, Madison Jewell, Jared Luckas, Emma Lesser, Tricia Nicholson, and Derrick Tang for XROMM data processing assistance.

## AUTHOR CONTRIBUTIONS

Conceptualization, Methodology, J.D.L-C and F.I.A-M.; Analysis & Software, J.D.L-C; Investigation, J.D.L-C and A.R.M.; Writing – Original Draft, J.D.L-C; Writing – Review & Editing, All; Funding Acquisition, F.I.A-M. (PI), N.G.H., and C.F.R.; Supervision, F.I.A-M. and C.F.R.

---

**Supplemental Figure 1.**
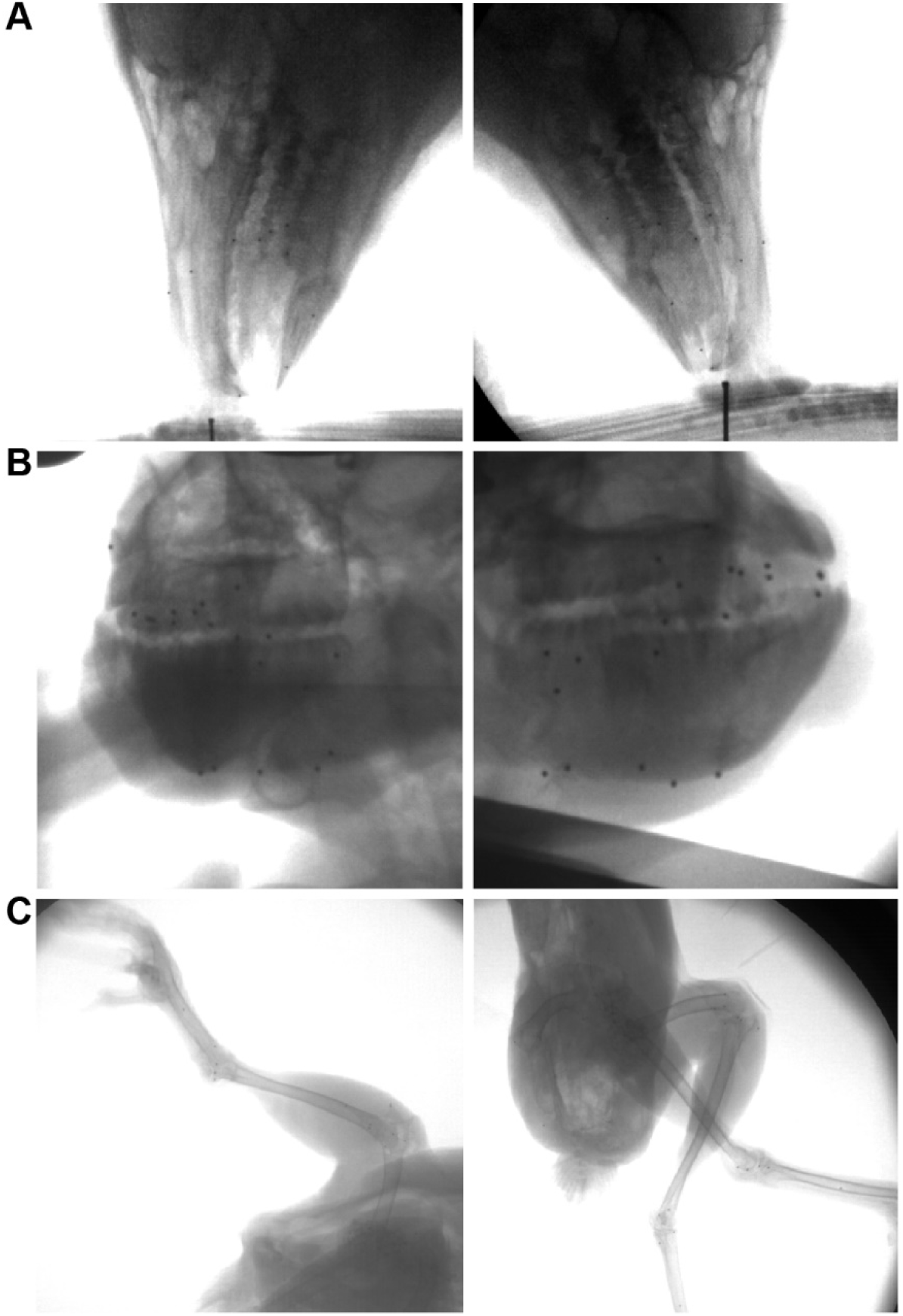
Example bi-planar X-ray images from each dataset. (A) Pig feeding; (B) Monkey feeding; (C) Bird range of motion. Note differences in marker number, size, and general appearance.

**Supplemental Figure 2.**
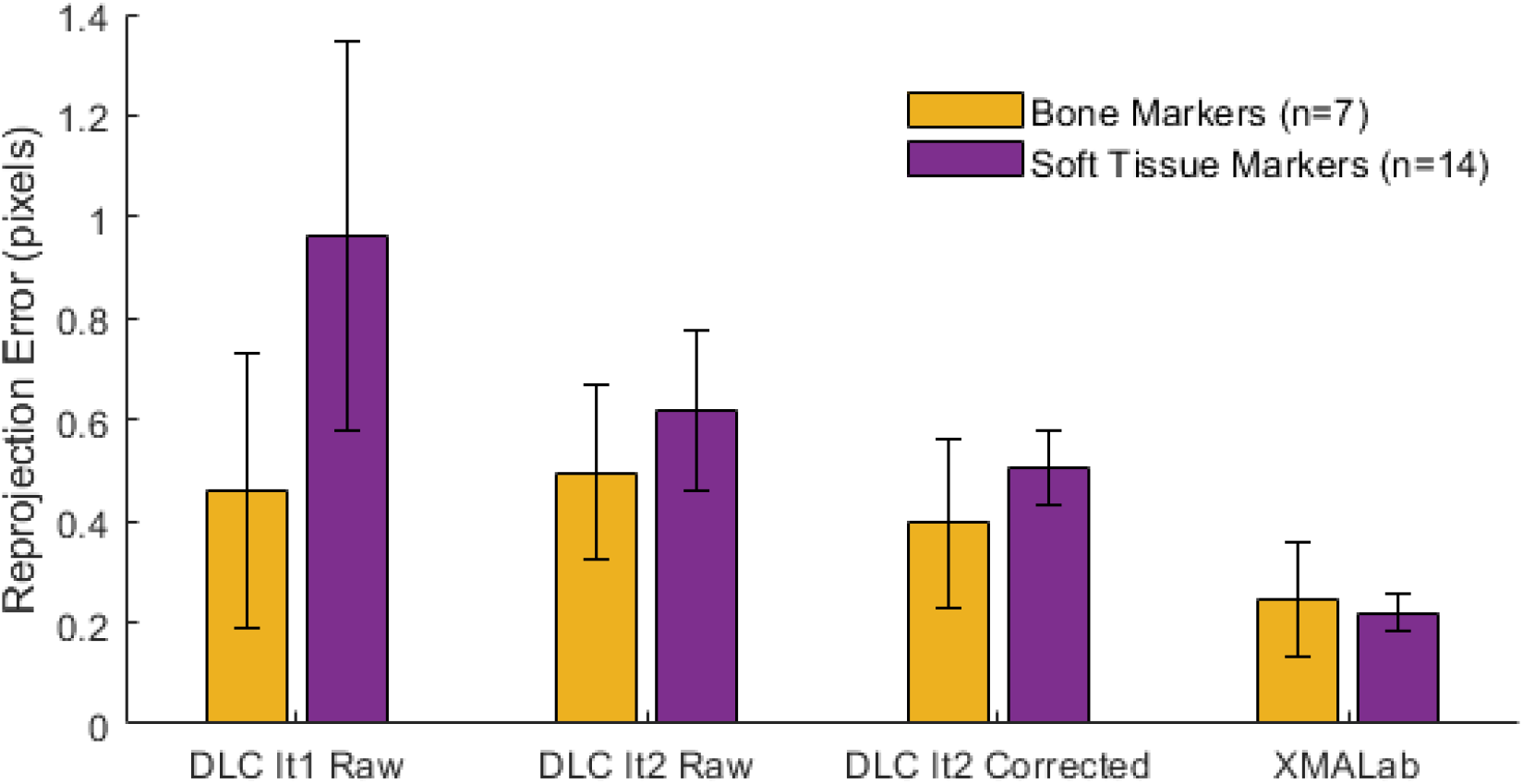
Comparison of mean marker reprojection errors across DeepLabCut network iterations and tracking mode. Error bars represent standard deviation. Note that the highest reprojection errors with the largest variance between markers occur in raw output of DeepLabCut’s first network iteration. Additionally, this is where the largest difference between bone and soft tissue markers is seen; the latter exhibit higher and more variable error. In the raw output of the second iteration— following augmentation of the training dataset with corrected frames—the mean reprojection errors are reduced, but the 3D positions do not yet reach the criterion for acceptable performance (not shown; see Methods). Manual correction of high-error frames is needed to reach this threshold. When the threshold is reached (DLC It2 Corrected) the reprojection errors are still, on average, higher than points tracked in XMALab alone. All values are the mean of reprojection error across all frames of the same test trial for each marker.

## REFERENCES

Bhullar, B. A. S., Manafzadeh, A. R., Miyamae, J. A., Hoffman, E. A., Brainerd, E. L., Musinsky, C. and Crompton, A. W. (2019). Rolling of the jaw is essential for mammalian chewing and tribosphenic molar function. Nature 566, 528–532.

Brainerd, E. L., Baier, D. B., Gatesy, S. M., Hedrick, T. L., Metzger, K. A., Gilbert, S. L. and Crisco, J. J. (2010). X-ray reconstruction of moving morphology (XROMM): precision, accuracy and applications in comparative biomechanics research. J. Exp. Zool. A. Ecol. Genet. Physiol. 313, 262–279.

Brust, C. A., Käding, C. and Denzler, J. (2019). Active learning for deep object detection. In VISIGRAPP 2019 - Proceedings of the 14th International Joint Conference on Computer Vision, Imaging and Computer Graphics Theory and Applications,.

Gintof, C., Konow, N., Ross, C. F. and Sanford, C. P. J. (2010). Rhythmic chewing with oral jaws in teleost fishes: a comparison with amniotes. J. Exp. Biol. 213, 1868–1875.

Granatosky, M. C., McElroy, E. J., Laird, M. F., Iriarte-Diaz, J., Reilly, S. M., Taylor, A. B. and Ross, C. F. (2019). Joint angular excursions during cyclical behaviors differ between tetrapod feeding and locomotor systems. J. Exp. Biol. 222.

Graving, J. M., Chae, D., Naik, H., Li, L., Koger, B., Costelloe, B. R. and Couzin, I. D. (2019). DeepPoseKit, a software toolkit for fast and robust animal pose estimation using deep learning. Elife.

Grood, E. S. and Suntay, W. J. (1983). A joint coordinate system for the clinical description of three-dimensional motions: Application to the knee. J. Biomech. Eng. 105, 136–144.

Hedrick, T. L. (2008). Software techniques for two-and three-dimensional kinematic measurements of biological and biomimetic systems. Bioinspiration and Biomimetics.

Insafutdinov, E., Pishchulin, L., Andres, B., Andriluka, M. and Schiele, B. (2016). Deepercut: A deeper, stronger, and faster multi-person pose estimation model. In Lecture Notes in Computer Science (including subseries Lecture Notes in Artificial Intelligence and Lecture Notes in Bioinformatics).

Iriarte-Diaz, J., Terhune, C. E., Taylor, A. B. and Ross, C. F. (2017). Functional correlates of the position of the axis of rotation of the mandible during chewing in non-human primates. Zoology.

Kambic, R. E., Roberts, T. J. and Gatesy, S. M. (2017). 3-D range of motion envelopes reveal interacting degrees of freedom in avian hind limb joints. J. Anat.

Knörlein, B. J., Baier, D. B., Gatesy, S. M., Laurence-Chasen, J. D. and Brainerd, E. L. (2016). Validation of XMALab software for marker-based XROMM. J. Exp. Biol. 219, 3701–3711.

Labuguen, R., Bardeloza, D. K., Negrete, S. B., Matsumoto, J., Inoue, K. and Shibata, T. (2019). Primate Markerless Pose Estimation and Movement Analysis Using DeepLabCut.

Manafzadeh, A. R. and Padian, K. (2018). ROM mapping of ligamentous constraints on avian hip mobility: implications for extinct ornithodirans. Proc. R. Soc. B 285, 20180727.

Martinez, C. M., McGee, M. D., Borstein, S. R. and Wainwright, P. C. (2018). Feeding ecology underlies the evolution of cichlid jaw mobility. Evolution (N. Y). 72, 1645–1655.

Mathis, M. W. and Mathis, A. (2019). Deep learning tools for the measurement of animal behavior in neuroscience.

Mathis, A., Mamidanna, P., Cury, K. M., Abe, T., Murthy, V. N., Mathis, M. W. and Bethge, M. (2018). DeepLabCut: markerless pose estimation of user-defined body parts with deep learning. Nat. Neurosci. 21, 1281–1289.

Menegaz, R. A., Baier, D. B., Metzger, K. A., Herring, S. W. and Brainerd, E. L. (2015). XROMM analysis of tooth occlusion and temporomandibular joint kinematics during feeding in juvenile miniature pigs. J. Exp. Biol. 218, 2573–2584.

Nath, T., Mathis, A., Chen, A. C., Patel, A., Bethge, M. and Mathis, M. W. (2018). Using DeepLabCut for 3D markerless pose estimation across species and behaviors. bioRxiv 14, 476531.

Orsbon, C. P., Gidmark, N. J. and Ross, C. F. (2018). Dynamic Musculoskeletal Functional Morphology: Integrating diceCT and XROMM. Anat. Rec. 301, 378–406.

Owen, S. F., Liu, M. H. and Kreitzer, A. C. (2019). Thermal constraints on in vivo optogenetic manipulations. Nat. Neurosci.

Parmiani, P., Lucchetti, C., Bonifazzi, C. and Franchi, G. (2019). A kinematic study of skilled reaching movement in rat. J. Neurosci. Methods.

Pereira, T. D., Aldarondo, D. E., Willmore, L., Kislin, M., Wang, S. S. H., Murthy, M. and Shaevitz, J. W. (2019). Fast animal pose estimation using deep neural networks. Nat. Methods.

Stringer, C., Pachitariu, M., Steinmetz, N., Reddy, C. B., Carandini, M. and Harris, K. D. (2019). Spontaneous behaviors drive multidimensional, brainwide activity. Science.

Tan, C., Sun, F., Kong, T., Zhang, W., Yang, C. and Liu, C. (2018). A survey on deep transfer learning. In Lecture Notes in Computer Science (including subseries Lecture Notes in Artificial Intelligence and Lecture Notes in Bioinformatics).

